# Aldehyde-based cryopreservation of whole brains

**DOI:** 10.64898/2026.03.02.708967

**Authors:** Macy Garrood, Alicia Keberle, Andria Slaughter, Allison Sowa, Emma L. Thorn, Claudia De Sanctis, Kurt Farrell, John F. Crary, Andrew T. McKenzie

**Affiliations:** Apex Neuroscience, Salem, OR 97317, USA; Microscopy and Advanced Bioimaging Core, Icahn School of Medicine at Mount Sinai, New York, New York, USA; Friedman Brain Institute, Departments of Pathology, Neuroscience, and Artificial Intelligence & Human Health, Icahn School of Medicine at Mount Sinai, New York, New York, USA; Neuropathology Brain Bank & Research Core and Ronald M. Loeb Center for Alzheimer’s Disease, Icahn School of Medicine at Mount Sinai, New York, New York, USA

**Author notes:** Corresponding author: Andrew T. McKenzie, MD, PhD.

**Keywords:** Brain banking, Cryopreservation, Aldehyde fixation, Ultrastructural quality, Biobanking

## Abstract

Long-term storage of aldehyde-fixed brain tissue is commonly performed in the fluid state. This has the potential to maintain morphology for many decades, but has been found to cause progressive loss of antigenicity over time for some biomolecules. While cryoprotection and subzero storage has been successfully used for brain tissue sections or blocks, methods for preserving whole brains using this approach have not been widely characterized. Here we present a protocol for the preservation of fixed whole brains using graded immersion cryoprotection and subzero temperature storage, which is one type of a more general approach that we refer to as aldehyde-based cryopreservation (ABC). Our method uses a gradual ramp-up of the osmotic concentration of cryoprotectants, leading to a final solution containing 50% (v/v) ethylene glycol and 30% (w/v) sucrose. We used CT imaging to track cryoprotectant penetration, finding that with the use of our protocol, approximately 10 months is required to reach equilibration throughout whole human brains. In our initial histological validation, we found that insufficient equilibration time prior to freezing led to apparent ice crystal artifacts seen on ultrastructural imaging of the white matter. After refining the protocol to allow adequate diffusion time, histologic data at both the light and electron microscopic levels showed preserved cellular architecture and ultrastructure after the process of cryoprotectant loading, freezer storage, and unloading. This protocol can be implemented using laboratory freezers or freezer rooms and provides a degree of resilience against freezer failures because the morphology of the fixed tissue is expected to remain preserved long-term in the fluid state even if rewarmed. Our approach may be valuable for laboratories seeking to enhance the long-term preservation of antigenicity in large brain tissue specimens for future research applications.

## Introduction

Many thousands of human brains are banked around the world yearly with the goal of future research study. Each banked brain is an irreplaceable specimen, often containing rare pathologies, unique genetic variants, or specific disease states that cannot be replicated. Additionally, many animal model specimens, such as those from long-term studies or unique experimental conditions, are similarly irreplaceable and may require robust long-term preservation. For this reason, it is critical to develop preservation methods that can maintain both structural and molecular features of brain tissue over extended time periods. Fluid preservation of aldehyde-fixed brain tissue at room or refrigerator temperature is a widely used method, due to its simplicity and ability to maintain morphological features for years or decades (1). However, studies have shown progressive loss of antigenicity during prolonged storage, as well as other potential biochemical changes, which could impact some types of future research applications. Although cryopreservation has proven successful for preserving molecular aspects of brain tissue sections or blocks, this technique can cause damage to cellular structures, in part due to the formation of ice crystals during freezing (2). As a result, there is a critical need to develop improved methods for long-term preservation that can both maintain structural integrity and minimize biomolecular alterations of whole brain specimens.

One alternative approach is to combine aldehyde fixation and cryopreservation. After the initial aldehyde fixation, the tissue can be immersed in cryoprotective agents and then cryopreserved. This approach prevents ice damage and the resulting morphological damage from unprotected cryopreservation, while also minimizing the loss of antigenicity associated with long-term storage in aldehyde solutions. We use the term aldehyde-based cryopreservation (ABC) to describe this general strategy of first applying aldehyde fixation and then performing cryoprotectant loading and subzero storage. Numerous previous studies have used this approach on sections or blocks of fixed brain tissue, finding that storage at -20°C or -25°C in cryoprotectant solutions can maintain both tissue morphology and protein immunoreactivity for extended periods (3–11). There have also been studies using this method on large segments of brain tissue or whole brains, using immersion to introduce the cryoprotective agents (12–15). However, there are still questions that are not completely resolved regarding ABC. One question is the extent to which any given protocol for immersion cryoprotection may cause structural alterations as visualized by microscopy. Although several studies have tested or employed these protocols and not identified significant alterations, replication of this finding would be useful. Another question is how long it takes for the solution to diffuse into the brain tissue, which determines how long of a waiting period is necessary before moving the brain to subzero storage.

To address these open questions, we describe a protocol for whole-brain cryoprotectant immersion following aldehyde fixation, enabling long-term storage at freezer temperatures. We then present two validation studies. First, using CT imaging of whole human brains, we characterize the time course of cryoprotectant penetration throughout the brain volume, providing a measure for when the tissue has reached sufficient equilibration for transfer to subzero storage. Second, through histological analysis of biopsy samples, we assess whether the cryoprotectant loading, freezer storage, and unloading steps introduce microscopic alterations. Our findings may be useful for investigators seeking to preserve intact brains for future research applications.

## Materials and Methods

### Anatomical donation procedures

Human whole body donations were performed by a partner organization operating under Oregon Health Authority regulations. Dog brains were obtained after euthanasia by a licensed veterinarian, with owner consent for research use. Pig brains were sourced as animal byproducts from regulated agricultural facilities. In both cases, animal tissue was obtained postmortem from animals that died for reasons unrelated to this study. The Apex Neuroscience Brain and Tissue Bank operates under an exemption determination issued by the Pearl Institutional Review Board.

### Initial tissue fixation

Human and pig brains were removed from the skull following standard procedures and immersion fixed in 10% neutral buffered formalin (NBF; Azer Scientific NBF55G) at 4°C for at least one month prior to further processing (**Table 1**) (16,17). The dog brains were perfusion fixed *in situ* with 10% NBF: donor #65 via transcardial cannulation approximately 90 minutes after euthanasia, and donor #177 via bilateral carotid artery injection approximately 20 minutes after euthanasia. Both were subsequently immersion fixed in 10% NBF at 4°C for at least one month prior to further processing.

**Table 1.** Reagents used in this study. MW: Molecular Weight.

| Chemical | Supplier | Identifier |
| --- | --- | --- |
| 10% Neutral Buffered Formalin | Azer Scientific | NBF55G |
| Glutaraldehyde | Thermo Fisher | A10500.0E |
| Ethylene glycol | CQ Concepts | r-671-5 |
| Sucrose | Lab Alley | SUCGA-500G |
| Polyvinylpyrrolidone (Average MW 40,000) | Sigma | PVP40-100G |

### Cryoprotectant penetration kinetics study

Three intact human brains were used to characterize the penetration kinetics of the cryoprotectant: a 71-year-old woman (donor #54, postmortem interval (PMI) of 117 hours), a 90+-year-old man (donor #137, PMI 47 hours), and a 75-year-old man (donor #180, PMI 46 hours). These represented a convenience sample of brains donated to the Apex Neuroscience Brain and Tissue Bank, with no specific exclusion criteria.

After fixation in 10% NBF for at least one month, whole brains were transferred through a series of cryoprotectant solutions. The brains were maintained at the temperature of 4°C, except for one week when they were left at room temperature. Step 1 was 10% ethylene glycol (v/v) added to the solution of 10% NBF. Step 2 was 30% ethylene glycol (v/v) added to 10% NBF. Step 3 (the final solution) was 50% ethylene glycol (v/v), 30% sucrose (w/v), and 1% PVP-40 (w/v), with the remainder of the solution being 10% NBF. Each intermediate step was maintained for at least one month before proceeding to the next concentration. The brains remained in the final solution until equilibration was confirmed by CT imaging.

CT scans were performed using the OmniTom Elite (Neurologica, Danvers, MA), a 16-slice portable CT scanner. Images were viewed with the Osimis Web Viewer. Baseline scans were obtained after fixation, with sequential scans after each cryoprotectant step. Equilibration was determined when the Hounsfield Unit values stabilized throughout the brain.

To assess macroscopic dimensional changes during cryoprotectant loading, we measured the brain width on coronal CT images at each imaging time point. For each scan, we identified the most anterior coronal slice in which the frontal horns of the lateral ventricles were clearly visible. The maximum transverse brain width was then measured at this level using the measurement tool in Orthanc Explorer 2. The frontal horns were chosen as the reference landmark as they are well-defined structures that are identifiable across serial scans.

### Histological validation study

Biopsy samples (approximately 1.5 x 1.5 x 1 cm) from two perfusion-fixed dog brains were used to assess whether cryoprotectant loading, freezer storage, and unloading introduce histological alterations. We performed an initial experiment (donor #65) and, based on those results, developed a refined protocol that was tested in a second experiment (donor #177).

### Initial protocol (donor #65)

Samples were collected from the sensorimotor cortex, thalamus, and subcortical white matter. For each region, samples were divided into cryopreserved and non-cryopreserved groups. Non-cryopreserved control samples were maintained in 10% NBF at 4°C during the cryoprotectant ramp up and down.

Cryopreserved samples were transferred through cryoprotectant solutions at 4°C. On day 1, samples were placed in 10% ethylene glycol (v/v) in 10% NBF. On day 2, samples were transferred to 30% ethylene glycol (v/v) in 10% NBF. On day 3, samples were transferred to the final solution: 50% ethylene glycol (v/v), 30% sucrose (w/v), and 1% PVP-40 (w/v), with the remainder being 10% NBF. After 3 days for equilibration, samples were transferred to −20°C for 3 days.

For cryoprotectant unloading, samples were rewarmed at 4°C for 24 hours, then transferred through decreasing concentrations: 30% ethylene glycol (v/v) in 10% NBF for 5 days, 10% ethylene glycol (v/v) in 10% NBF for 7 days, and finally 10% NBF alone. At the end of this, glutaraldehyde (2% v/v) was added to both cryopreserved and control samples for one day prior to processing for electron microscopy.

### Refined protocol (donor #177)

Based on the ultrastructural artifacts we observed in white matter samples from the initial protocol, we developed a refined protocol with three main modifications: (1) a slower loading protocol with more time for equilibration prior to moving the samples to the freezer, (2) a more gradual unloading protocol with smaller concentration decrements, and (3) inclusion of glutaraldehyde earlier in the loading and unloading steps.

Samples were collected from parietal and temporal cortex and nearby subcortical white matter. For each region, samples were divided into cryopreserved and non-cryopreserved groups. Non-cryopreserved control samples were maintained in 10% NBF with 2% glutaraldehyde (v/v) at 4°C, to match the glutaraldehyde exposure of the cryopreserved group.

Cryopreserved samples were transferred through cryoprotectant solutions at 4°C. On day 1, samples were placed in 10% ethylene glycol (v/v) in 10% NBF. On day 5, samples were transferred to 30% ethylene glycol (v/v) in 10% NBF, with 2% glutaraldehyde (v/v). On day 9, samples were transferred to the final solution: 50% ethylene glycol (v/v), 30% sucrose (w/v), 2% glutaraldehyde (v/v), 10% (v/v) of 10% NBF, with the remainder consisting of 0.01M PBS. After 3 days for equilibration, the samples were transferred to −20°C for 7 days.

For cryoprotectant unloading, samples were rewarmed at 4°C for 24 hours, then transferred through a gradual 18-day protocol with decreasing concentrations of ethylene glycol and sucrose (**Table 2**). Glutaraldehyde (2% v/v) was maintained throughout all unloading steps. Following unloading, samples remained in 10% NBF with 2% glutaraldehyde at 4°C for several months until processing for electron microscopy.

**Table 2.** Cryoprotectant unloading steps for the refined protocol (donor #177). Glut: Glutaraldehyde. PBS: Phosphate buffered saline. NBF: Neutral buffered formalin.

| Day | Ethylene glycol (v/v) | Sucrose (w/v) | 0.01M PBS (v/v) | 10% NBF (v/v) | Glut. (v/v) |
| --- | --- | --- | --- | --- | --- |
| 1 | 40% | 25% | 30% | 28% | 2% |
| 4 | 30% | 20% | 23% | 45% | 2% |
| 7 | 20% | 15% | 18% | 60% | 2% |
| 11 | 10% | 10% | 13% | 75% | 2% |
| 15 | 5% | 5% | 8% | 85% | 2% |
| 18 | 0% | 0% | 0% | 98% | 2% |

### Histological methods

For light microscopy, brain tissue was placed into cassettes for processing and embedded in paraffin. Paraffin-embedded brain sections 6 μm thick were baked, deparaffinized, and stained for Hematoxylin and Eosin (H&E). Digital images of the stained sections were captured at 40X as whole slide images (WSIs) using an Aperio GT450 high-resolution scanner (Leica Biosystems).

For electron microscopy on the initial protocol, we followed a previously described protocol (18). Briefly, the samples were cut into 1 mm^3^ pieces and placed in mPrep/s capsules. Using an ASP-2000, the samples were rinsed in 0.1M sodium cacodylate buffer, stained with 2% osmium tetroxide in 0.1M sodium cacodylate, then 2.5% w/v potassium ferricyanide in 0.1M sodium cacodylate, rinsed in water, immersed in 1% w/v aqueous thiocarbohydrazide, rinsed in water, stained with 2% aqueous osmium tetroxide, rinsed in water, stained with 1% uranyl acetate, rinsed in water, and finally stained with lead aspartate. After rinsing in water, the samples were dehydrated in an ascending acetone series, infiltrated with resin overnight, and polymerized in a 60C oven. Once polymerized, the samples were sectioned at 70nm thickness. Sections were placed on a grid, stained with uranyl acetate and lead citrate, and imaged at 80kV in an FEI Tecnai T12 transmission electron microscope equipped with an AMT Nanosprint12 camera.

For electron microscopy on the refined protocol, tissue samples were processed as previously described (19). Briefly, the tissue was processed using an adapted NCMIR protocol for enhanced contrast. This included sequential treatments with tannic acid, reduced osmium, thiocarbohydrazide, osmium, and uranyl acetate at room temperature, followed by lead aspartate staining at 60°C. Samples were dehydrated through graded ethanol, infiltrated with Embed 812 epoxy resin (EMS), and polymerized for 72 hours at 60°C. Ultrathin sections (70 nm) were cut using a Leica UC7 ultramicrotome and collected on nickel slot grids. Images were acquired on a HT7500 transmission electron microscope (Hitachi High-Technologies, Tokyo, Japan) using an AMT NanoSprint12 12-megapixel CMOS TEM Camera System, with minimal contrast adjustments applied during acquisition.

Two raters (M.G. and A.S.), blinded to experimental condition, independently scored each electron micrograph of the refined protocol for ultrastructural preservation quality using a subjective 5-point scale (1 = best preservation, 5 = worst preservation). The interrater reliability for these grades was calculated using the intraclass correlation coefficient (ICC), applying a model with agreement estimation, single unit of analysis, and two-way random-effects. The ICC values were interpreted using previously established guidelines (20). For analysis, rater scores were averaged (arithmetic mean) to produce a single score per image. These image-level scores were then averaged within each sample to yield one arithmetic mean score per sample. Differences between the cryopreserved and non-cryopreserved groups were assessed using the Wilcoxon rank-sum test due to the small sample size (n = 6 per group).

## Results

### Choice of cryoprotective solution

A wide variety of cryoprotective agents (CPAs) or mixtures could potentially prevent ice crystal formation at a storage temperature of -20°C. Our goal was to identify a mixture that would not only prevent ice formation but also serve well to maintain tissue structure as a fluid preservative, especially in the case of a freezer failure. We initially tested glycerol, which we reasoned would be desirable due to its long established history as a fluid preservation agent and its high viscosity (1). However, we found that pig brain tissue exposed to a concentration of 90% (v/v) glycerol for two days at 4°C demonstrated significant surface browning. This phenomenon of glycerol-induced browning has been previously reported numerous times, for example in the context of embalming (21) and tissue clearing (22). It is likely due to the Maillard reaction. While this browning does likely not impact a significant number of downstream applications, it could interfere with some analyses and was therefore deemed undesirable. This observation led us to explore alternative CPA additives that did not contain glycerol. We used ethylene glycol (50% v/v), sucrose (30% w/v), and PVP-40 (1% w/v), building on a previously designed preservation method that performed storage at -20°C (4). We tested this storage method on a pig brain, finding that it did not cause rapid surface browning nor any obvious volumetric changes on gross visual examination. Furthermore, we found that CT scans could be used to track the increases in the concentration of ethylene glycol.

### Penetration rate and effect on volumes in human brains

For the first two cryoprotectant loading steps (10% and 30% ethylene glycol), we waited approximately one month between each concentration increase, which our CT scans revealed was not sufficient for full equilibration throughout the entire brain volume (**Fig 1**). However, we reasoned that brain-wide equilibration was not necessary at these intermediate steps, as it is only the outermost brain tissue that would be exposed to the higher concentration and thus the largest osmotic concentration difference. The more inner parts of the brain tissue would instead gradually increase in cryoprotectant concentration over time through diffusion, naturally preventing a large increase in osmotic concentration at any specific region. This approach allowed us to substantially decrease the waiting period before transferring the brain to subzero temperatures, making the protocol more feasible for implementation in brain banking facilities. For the final concentration step, we did wait for full equilibration as confirmed by CT imaging, with Hounsfield Unit values stabilizing between outer cortical and deep brain regions after 276 days. Notably, we found that white matter appeared to have substantially slower cryoprotectant diffusion kinetics than grey matter.

**Fig 1.**
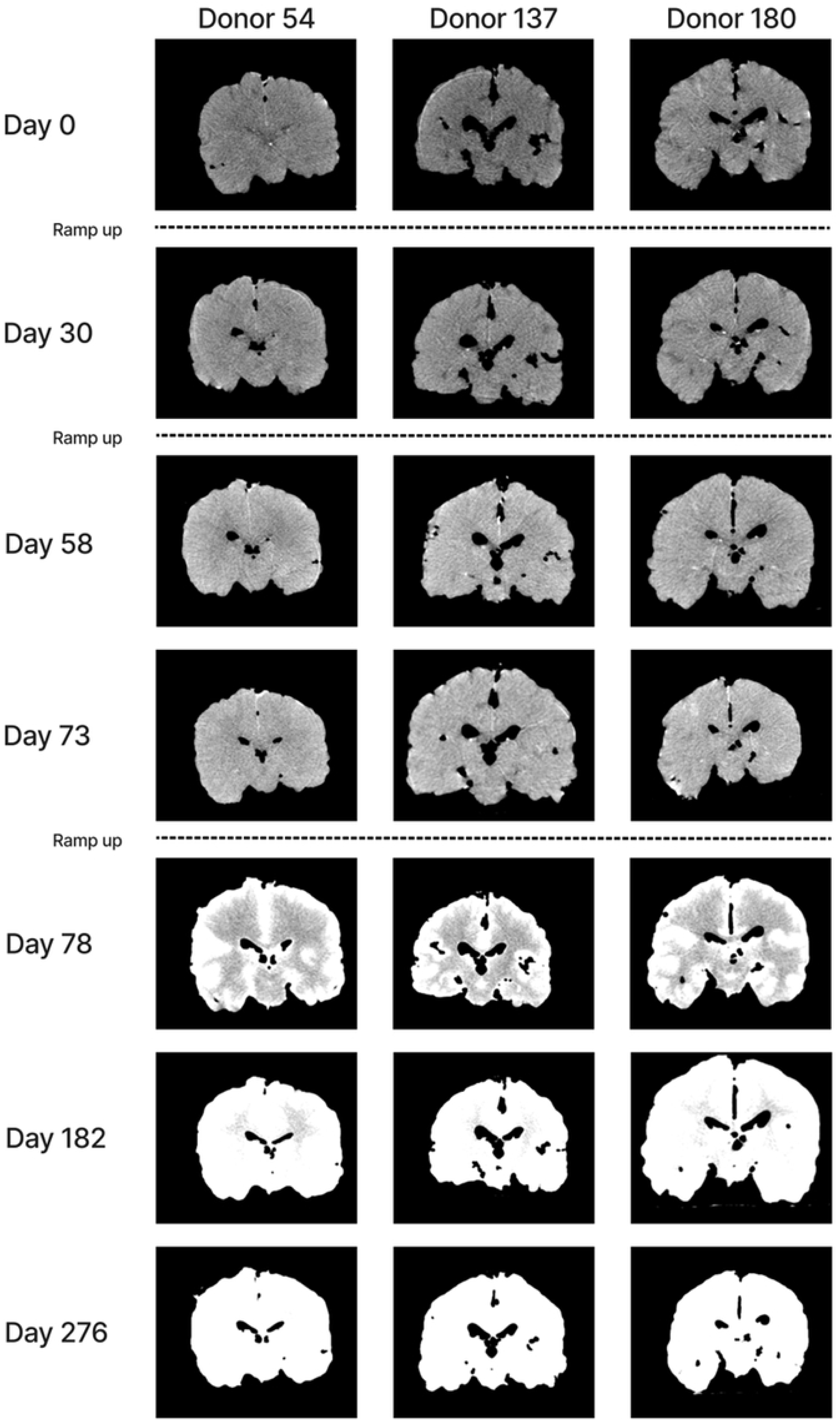
CT imaging of cryoprotectant penetration in whole human brains. Coronal CT sections from three donors (donor #54: 71-year-old female, postmortem interval (PMI) 117 hr; #137: 90+-year-old male, PMI 47 hr; #180: 75-year-old male, PMI 46 hr) during stepwise cryoprotectant loading. Dashed lines indicate the concentration increases: 10% ethylene glycol (added on day 0), 30% ethylene glycol (added on day 30), and final solution containing 50% ethylene glycol, 30% sucrose, and 1% PVP-40 (added on day 73). Increasing brightness reflects rising Hounsfield Unit values as cryoprotectant diffuses inward. Note that grey matter equilibrates faster than white matter, with deep white matter regions showing delayed penetration. By day 276, the uniform signal intensity indicates equilibration throughout all of the brain.

To assess for macroscopic dimension changes during cryoprotectant loading, we measured the maximum transverse brain width at the level of the frontal horns of the lateral ventricles on serial CT images. For donors #54 and #137, the brain width remained stable throughout the loading protocol, with no appreciable volumetric changes observed across all time points (**S1 Data**). Donor #180 similarly showed stable measurements for the majority of the time points, but then at one time point exhibited an abrupt increase of approximately 10% in the measured width, which persisted stably across the final four time points (**S1 Data**). One possible explanation is that this could be a measurement artifact. In particular, as the cryoprotectant solution entered the ventricular spaces, the increased contrast within the ventricles may have altered the anteroposterior level at which the frontal horns were first visualized on coronal sections, shifting the measurement plane posteriorly to where the brain is naturally wider. However, we cannot rule out an alternative explanation or a true volume change, and future studies using more precise volumetric methods on more donor brains could help to clarify this.

After two months of freezer storage at -20°C, one of the brains was removed for examination. We found that there were no macroscopic signs of ice crystal formation. The brain was firm but not frozen, and was easily able to be sectioned into for extracting a biopsy sample. There was a degree of surface browning, which accumulated during the approximately 10-month cryoprotectant loading period at 4°C, though this was not as severe as that observed with 90% glycerol. The browning was primarily present on the surface, with a gradient in the outermost few millimeters of the tissue.

### Histological assessment of the initial protocol

In our initial experiment using tissue from donor #65, we assessed whether cryoprotectant loading, freezer storage at -20°C, and subsequent unloading would alter the tissue ultrastructure compared to non-cryopreserved controls maintained in fixative solution. We found that grey matter samples from the sensorimotor cortex and thalamus showed comparable preservation between cryopreserved and control conditions, with no differences in the structure of cellular membranes, and no obvious artifacts attributable to the cryopreservation process (**Fig 2**). Neither condition showed perfect ultrastructural preservation, but the alterations observed were consistent with expected postmortem changes rather than cryopreservation-induced damage (23,19).

**Fig 2.**
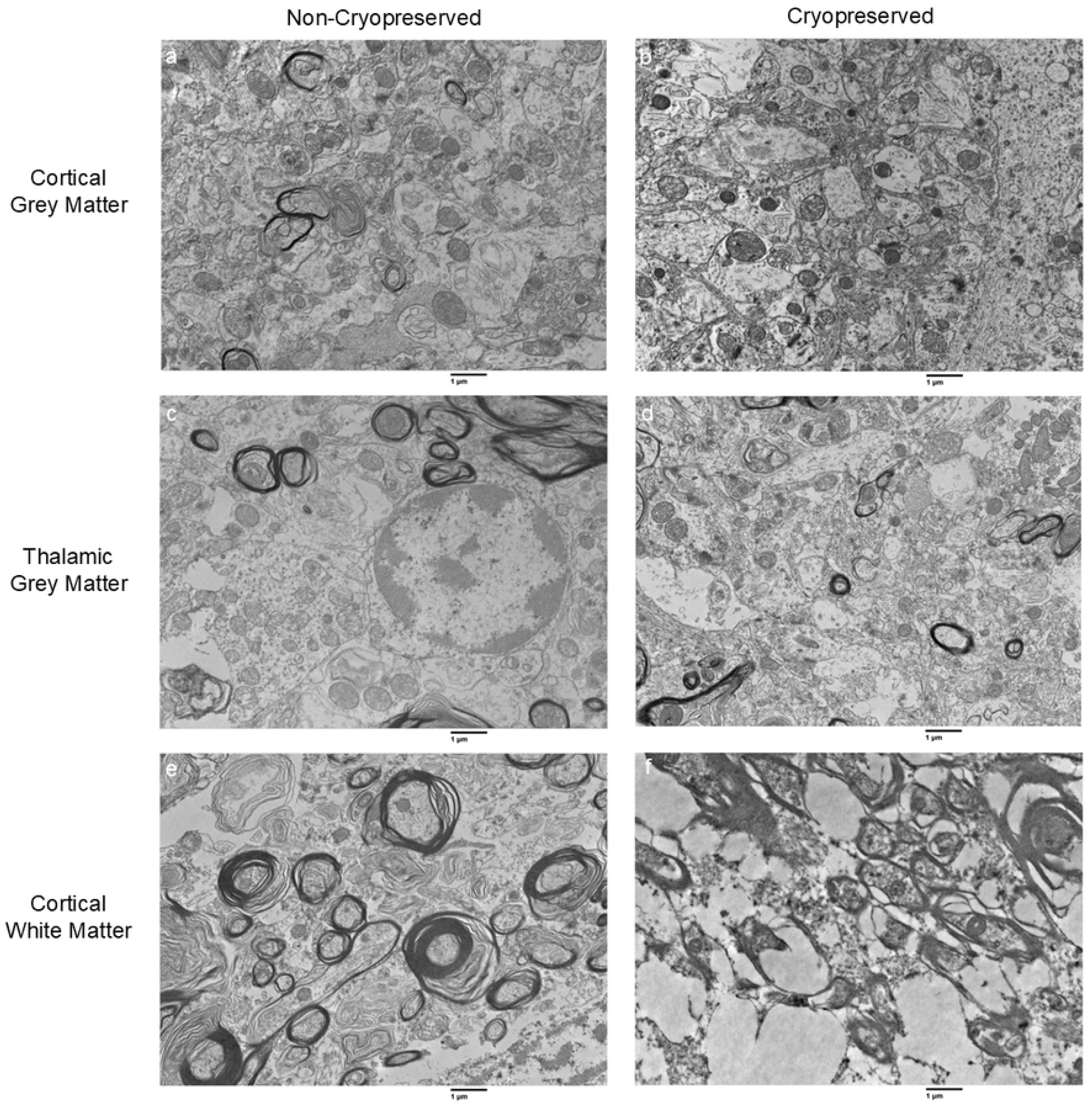
Electron microscopy of cryopreserved and non-cryopreserved brain tissue from the initial protocol. Left column: non-cryopreserved controls maintained in fixative solution prior to embedding for electron microscopy. Right column: tissue after cryoprotectant loading, storage at -20°C, and unloading. **a**, **b**: Cortical grey matter. **c**, **d**: Thalamic grey matter. **e**, **f**: Subcortical white matter. Grey matter shows comparable preservation between conditions. Cryopreserved white matter (**f**) exhibits large void spaces with compressed surrounding tissue, consistent with ice crystal formation due to incomplete cryoprotectant penetration prior to the freezer storage. Scale bars: 1 μm.

However, subcortical white matter samples showed a different outcome. While non-cryopreserved white matter displayed myelinated axon profiles with expected postmortem changes such as delamination, cryopreserved white matter exhibited prominent artifacts characterized by large, sharply delineated void spaces that displaced and compressed the surrounding tissue (**Fig 2**). These irregular vacuolar structures ranged from sub-micron to approximately 10 μm in diameter and were distributed throughout the tissue, with intervening regions of compressed myelinated fibers between them. The morphology of these artifacts – i.e., discrete empty spaces with sharp boundaries and compressed surrounding tissue – is consistent with ice crystal formation (2). We hypothesize that the 3-day cryoprotectant loading period was insufficient for complete equilibration in white matter, which has slower diffusion kinetics than grey matter. This would have left residual freezable water in white matter compartments, leading to ice formation during storage at -20°C. Ice crystal formation has previously been reported in fixed brain tissue when cryoprotectant infiltration is incomplete (12). It is also possible that insufficient glutaraldehyde fixation prior to EM processing or osmotic stress during cryoprotectant unloading contributed to these artifacts. However, the sharp morphology of the artifacts and their restriction to the slower-diffusing white matter suggest that ice crystal formation is the sole cause.

### Histological assessment of the refined protocol

We next evaluated tissue preservation using the refined cryopreservation protocol, which incorporated longer equilibration times, more gradual cryoprotectant unloading steps, and glutaraldehyde fixation. Light microscopy evaluation of both cryopreserved and non-cryopreserved samples using this protocol showed that the tissue architecture was preserved across all specimens (**Fig 3**). In both conditions, cell bodies, nuclei, and blood vessels remained clearly visible, and we observed no void spaces or tissue disruption patterns characteristic of ice crystal formation. This indicates that the cryoprotectant loading, freezer storage, and unloading protocol successfully maintained tissue integrity at the light microscopic level.

**Fig 3.**
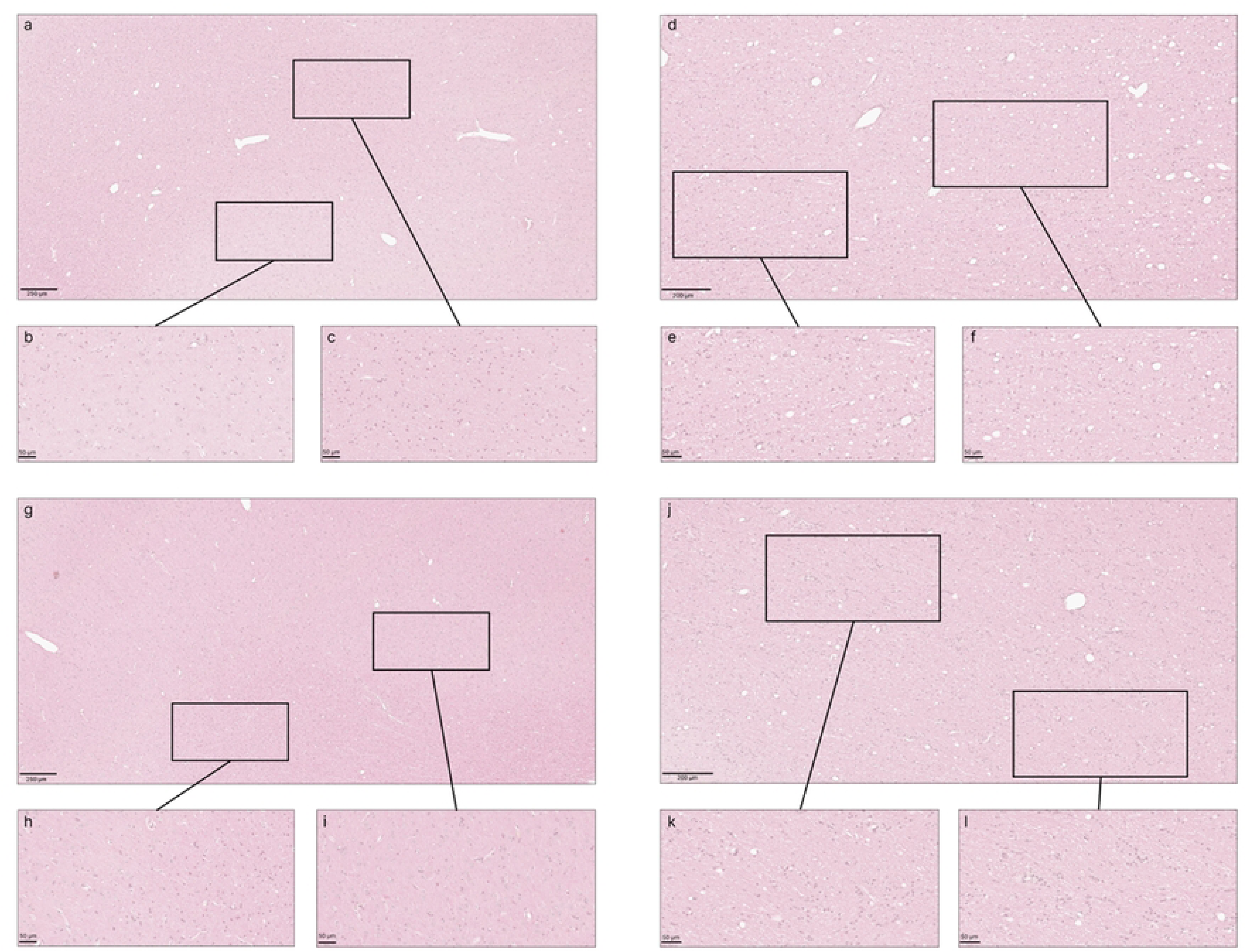
Representative light microscopy images comparing cryopreserved and non-cryopreserved canine brain tissue samples. H&E staining of cortical grey matter (**a-c**, **g-i**) and subcortical white matter (**d-f**, **j-l**). Top rows show cryopreserved samples (**a-c**: sample ID C1; g-i: sample ID C5) that underwent cryoprotectant loading, storage at -20 °C, and unloading. Bottom rows show non-cryopreserved controls (**d-f**: sample ID N1; **j-l**: sample ID N5) maintained in fixative throughout. Progressive magnification from overview to cellular detail shows that the tissue architecture is preserved in both conditions, with no observable differences in preservation quality between the cryopreserved and control samples in either grey matter or white matter. Scale bars: **a**, **d**: 250 μm; **g**, **j**: 200 μm; **b**, **c**, **e**, **f**, **h**, **i**, **k**, **l**: 50 μm.

Electron microscopy of grey matter and white matter samples showed comparable preservation between the cryopreserved samples and non-cryopreserved controls (**Fig 4**). As with the initial experiment, neither condition showed perfect ultrastructural preservation, with expected postmortem alterations. Qualitatively, we found that there was no clear effect of the cryopreservation protocol. For example, there were no large void spaces observed in the white matter that were attributed to ice crystal artifacts, as were seen in the initial protocol. In order to assess this, two blinded raters independently scored each image for ultrastructural preservation quality on a subjective 5-point scale (1 = best, 5 = worst). Blinded ratings of the preservation quality in electron microscopy images showed good interrater reliability (ICC = 0.76, 95% CI: 0.69 to 0.82). There was no significant difference in the mean preservation scores between the six cryopreserved samples (mean = 2.10, median = 2.12) and the six non-cryopreserved control samples (mean = 2.54, median = 2.63; Wilcoxon rank-sum test, W = 11, p = 0.30). These results suggest that the refined cryopreservation protocol successfully preserves the tissue throughout the cryoprotectant loading, freezer storage, and unloading process, without introducing detectable ultrastructural artifacts.

**Fig 4.**
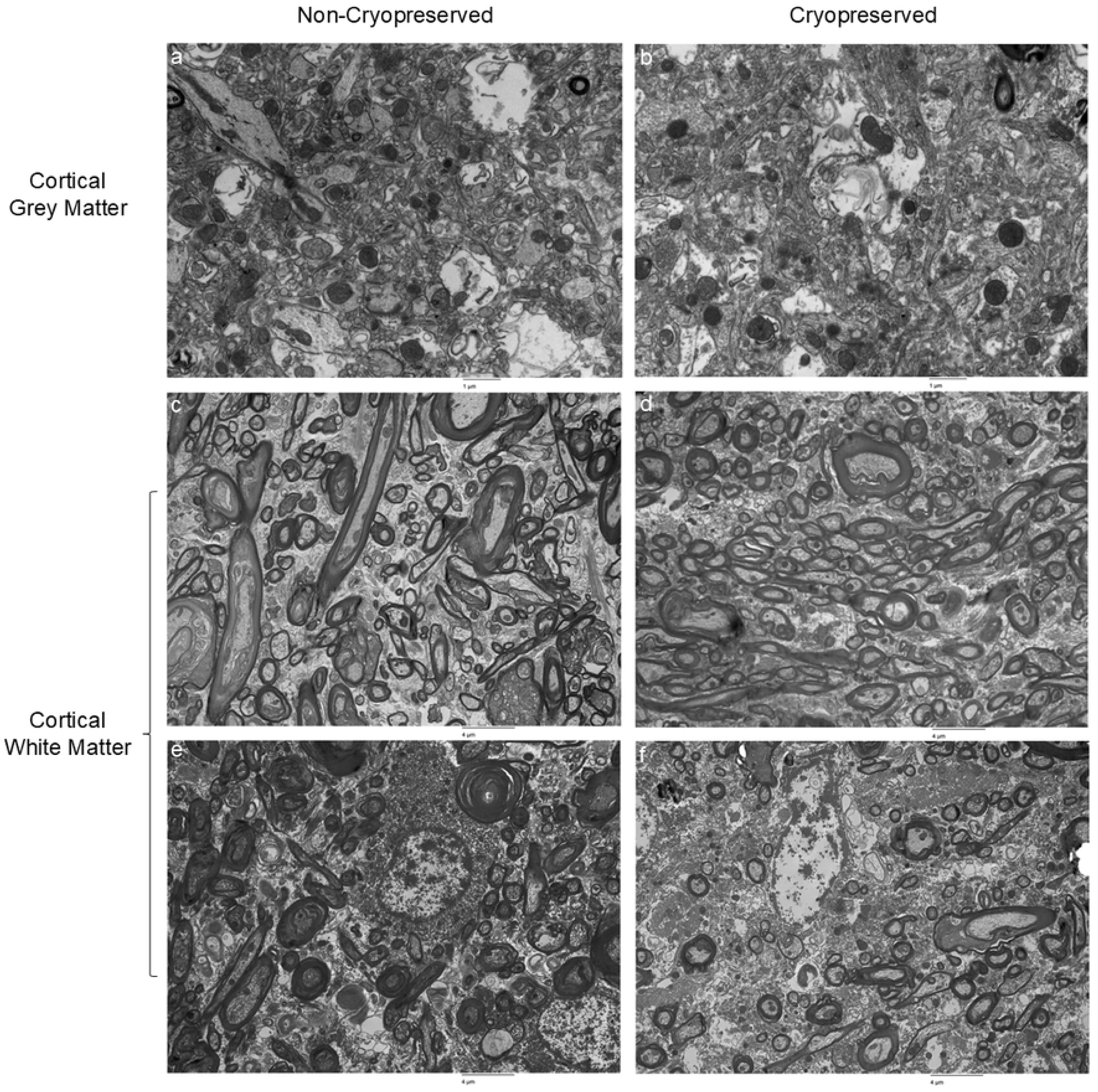
Representative electron micrographs of cryopreserved and non-cryopreserved brain tissue from the refined protocol. Left column: non-cryopreserved controls maintained in fixative solution prior to embedding for electron microscopy. Right column: tissue after cryoprotectant loading, storage at -20°C, rewarming, and cryoprotectant unloading. **a**, **b**: Cortical grey matter. **c**-**f**: Cortical white matter. Both grey matter and white matter show comparable preservation between cryopreserved and non-cryopreserved conditions, without the artifacts that were observed in the white matter in the initial protocol. Scale bars: **a**, **b**: 1 μm; **c**-**f**: 4 μm.

## Discussion

In this study, we describe our protocol for whole-brain cryoprotectant immersion following aldehyde fixation, enabling long-term storage at -20°C. This is one implementation of the ABC approach to long-term preservation. Using CT imaging, we characterized the time course of cryoprotectant penetration in whole human brains, finding that with this protocol, approximately 10 months is required for full equilibration of the final preservation solution. In our histology studies, we found that cryoprotectant loading, freezer storage, and unloading did not introduce significant alterations to tissue architecture or ultrastructure, as long as the time for cryoprotectant loading was adequate. This method offers several advantages for brain banking applications, as it can be implemented using standard laboratory freezers, it keeps the tissue easily accessible for sampling, and it relies upon widely available, well-characterized cryoprotective agents.

### Cryoprotectant formulation

A well-established protocol in the literature for long-term freezer storage of fixed brain tissue, Watson et al. (1986), uses a mixture of 30% ethylene glycol, 30% sucrose, and 1% polyvinylpyrrolidone (PVP-40) (4). Storage of fixed brain tissue under these conditions has been reported to maintain both tissue ultrastructure and antigenicity for multiple decades (4,8). We modified this protocol in several ways to optimize it for our purposes.

First, we increased the concentration of ethylene glycol from 30% (v/v) to 50% (v/v). At this concentration, ethylene glycol alone is sufficient to ensure the complete prevention of ice formation, because any solution with a melting point below the storage temperature cannot freeze (24). Raising the concentration of ethylene glycol further would lead to a risk of crystallization of ethylene glycol solutes.

Second, our final protocol uses fixative as a component of the solution. The rationale for this is twofold. First, we want to prevent the possibility of microbial growth during the approximately 10-month diffusion period at refrigerator temperature, which is much longer than required for tissue sections. We note that the residual fixative within the tissue after the initial fixation would likely be sufficient for this purpose, so adding fixative to the cryoprotectant loading solutions is a more conservative approach and may not be necessary. Second, we want the solution to function as an effective long-term fluid preservative in its own right. This way, if circumstances arise wherein freezer storage becomes not possible, the tissue would remain adequately preserved in the fluid state at room or refrigerator temperature for the long-term without the need for any solution changes.

Third, although PVP-40 was included in the original formulation (4), we removed it from our final protocol. We found it impractical for our whole brain application, as it was difficult to prepare large volumes of solution containing it, because PVP dissolves so slowly. Since 50% ethylene glycol alone provides sufficient cryoprotection at -20°C, and the sucrose component contributes to solution viscosity, we determined that PVP was not essential for our purposes.

We maintained the same concentration of sucrose at 30% w/v as in the original protocol (4). Sucrose will increase the viscosity of the solution, which is expected to slow the molecular motion of biomolecules and aid in long-term preservation in the fluid state (1). Consistent with this, sucrose has been found to be an effective fluid preservative for heart valves stored at 4°C (25). Additionally, ethylene glycol has been reported to be an effective agent for improving the preservation of fixed tissue in the fluid state (26,27).

### Loading and unloading of cryoprotectants

The extent to which gradual osmotic concentration changes are necessary when introducing cryoprotectants to aldehyde-fixed tissue is not yet entirely clear. Fixatives themselves are not thought to exert a substantial effective osmotic force, because they freely diffuse across cell membranes (28,29). Some studies have reported that fixed tissue can still have osmotic sensitivity, although the ones we identified employed relatively short fixation times, such as 4 hours or less (30,31). In contrast, one study found that rat brain tissue fixed with glutaraldehyde for 24 hours is entirely resistant to osmotic stress (32). Other studies have also found that an adequate degree of aldehyde fixation prior to cryoprotection prevents cellular alterations due to osmotic stress (33–35).

Several studies using similar cryoprotection protocols for fixed brain tissue have employed an immediate transfer to the final cryoprotectant concentration without a gradual ramp-up (8,11). For example, in the Watson et al. (1986) protocol, which is the primary inspiration for this method, cryoprotected fixed brain sections were placed immediately in the final cryoprotectant mixture (4). They reported that the ultrastructure remained normal after washing, although they did note transient shrinkage of the tissue upon cryoprotectant removal that resolved within 30 minutes in PBS. Other protocols have used a single intermediate cryoprotectant ramp up step (12,13).

In contrast, our final protocol uses three ramp-up steps (10% ethylene glycol, 30% ethylene glycol, then the final solution). For whole brain protocols such as ours, the stepwise gradient is likely only relevant to the outermost surface of the brain tissue. However, because we do not wait for equilibration until the final step, which takes by far the longest time, our use of the slower ramp-up approach only adds a relatively small amount of time compared to the overall protocol. Future studies could determine whether this degree of caution regarding rapid osmotic concentration changes is necessary, or whether more rapid loading and unloading would produce equivalent results with less time and complexity.

### Equilibration time

There is some previously reported data on how much time has been used for cryoprotectant immersion in brain tissue. One study reported that fixed monkey brain blocks equilibrated with 30% sucrose in 4-7 days, as determined by sinking (12). Another study reported that perfusion-fixed elephant brains equilibrated in their cryoprotectant solution (30% glycerol, 30% ethylene glycol) in approximately 3 weeks (13).

In this study, our CT data demonstrate that the full cryoprotectant loading protocol we used requires approximately 10 months at 4°C to reach a uniform distribution throughout intact human brains, with the final concentration ramp up and equilibration requiring approximately 8 months. Performing the equilibration at room temperature would accelerate diffusion, but we chose to use refrigerator temperature to minimize any potential biochemical changes during the loading period. Our CT imaging also revealed that white matter equilibrates substantially slower than grey matter. White matter consists of densely packed myelinated axons, and the lipid-rich myelin sheath may impede the diffusion of water-soluble cryoprotectants.

One important limitation of our use of CT imaging to assess the penetration rate is that it is a measure of the combined X-ray attenuation of all solutes. Our final solution contains both ethylene glycol (MW 62 g/mol) and sucrose (MW 342 g/mol), which will diffuse at different rates, with ethylene glycol diffusing faster due to its smaller size. Because our concentration of ethylene glycol alone provides sufficient cryoprotection at -20°C, and because it penetrates faster than sucrose, adequate ice prevention will still be assured by the time a uniform CT signal is reached, regardless of which solute contributes more to the CT signal.

Instead of using cryoprotectant immersion, another possibility would be to perfuse cryoprotectant immediately after perfusion fixation. This method can achieve a more rapid distribution of cryoprotectants throughout the brain when used at the time of death in ideal laboratory circumstances (35,36). However, in the context of postmortem human brain banking, perfusion quality is unreliable due to agonal factors, the postmortem interval, and vascular pathology in aged or diseased brains (37,38). In the cases in which perfusion quality is low, there will be osmotic damage due to the introduction of cryoprotectants without prior fixation, inadequate intracellular penetration of cryoprotectants in the absence of fixation, and/or ice damage in areas with poor perfusion (39,35,2). Therefore, we reasoned that the ABC approach, while substantially slower because it relies on cryoprotectant immersion, is more reliable for achieving structural preservation across the diverse range of specimens encountered in brain banking.

### Long-term preservation

A key limitation of this study is that we did not test long-term storage effects. However, our approach closely parallels other protocols that have been shown to maintain both tissue morphology and protein immunoreactivity for years (11) or decades (8). Theoretically, the Q10 temperature coefficient predicts that most chemical reactions will slow by a factor of approximately 2 for every 10°C decrease in temperature. However, empirical evidence suggests that preservation at subzero temperatures with cryoprotectants substantially exceeds these kinetic predictions. For example, one study examined unfixed cat brains cryoprotected with 15% glycerol and stored at -20°C for up to 7.25 years, finding that parts of the brain retained cellular structure (40,41). This contrasts sharply with unfixed brain tissue stored at 4°C, which generally shows rapid cellular degradation within days to weeks (23). As another example, fixed brain sections stored in cryoprotectant at -20°C have been found to maintain immunoreactivity for over 20 years, whereas before this method was developed, sections stored in standard buffers at refrigerator temperature could maintain this type of antigenicity for only a few days (8).

Samples naturally preserved in permafrost demonstrate that subzero temperatures even above -20°C (around -7 to -10°C) can enable very long-term preservation of some types of biological specimens. For example, DNA structure has been recovered from permafrost specimens over 1 million years old (42) and collagen ultrastructure has been found to be preserved for 40,000 years (43). Notably, our specimens are preserved in a viscous fluid state rather than being frozen, which differs mechanistically from permafrost. Still, taking the available evidence together, we expect that the combination of aldehyde crosslinking, cryoprotectants to increase solution viscosity, and storage at -20°C will maintain the morphomolecular integrity of brain tissue for many decades.

Another possibility with the use of the ABC approach is to store the tissue at a lower temperature, such as -40°C or -80°C. Whether lower temperatures would provide meaningful additional preservation benefit beyond -20°C is unclear, as -20°C storage with cryoprotectants already dramatically slows biochemical reactions relative to refrigerator temperatures, resulting in negligible changes in antigenicity for decades (8). However, some investigators may prefer storage at lower temperatures. Our 50% (v/v) ethylene glycol solution alone has a melting point of -36.8°C (24). The sucrose in the solution would depress the melting point further, but how much it would do so is, to the best of our knowledge, not well characterized. Therefore, modifying the cryoprotectant formulation may be necessary for the use of lower temperature storage. One option would be to increase the total concentration of permeating cryoprotectants, for example by incorporating DMSO. DMSO and glycerol mixtures have previously been used together for immersion cryoprotection of fixed brain tissue blocks prior to freezing and sectioning (12,5). Future studies could evaluate whether modified cryoprotectant formulations and lower temperature storage provide measurable benefits over the -20°C protocol described here.

Notably, standard fluid preservation of aldehyde-fixed brain tissue at room or refrigerator temperatures has also been found to maintain many morphological features for extended periods (1). The primary documented benefit of subzero storage in cryoprotectants is the prevention of progressive antigenicity loss that has been observed for some epitopes during prolonged fixative storage (4,8). The extent to which cryoprotection and subzero storage may provide additional preservation benefits beyond improved retention of antigenicity, such as reduced lipid oxidation, is an area for future investigation. Another consideration is that the required 10-month equilibration period entails a duration of fluid storage where some degree of antigen masking would already be expected to have occurred. However, because the decline in immunoreactivity for moderately fixation-sensitive antigens is a progressive process that continues to worsen even after the first year (1), stabilizing the tissue state at 10 months is thought to still offer a long-term advantage over indefinite storage at higher temperatures.

### Limitations

This study has several additional limitations. First, our protocol evolved during the study, with the refined histological validation protocol incorporating changes prompted in part due to the artifacts we observed in the initial experiment. Second, our sample sizes were small. Third, we used donated dog tissue because it allowed for shorter postmortem intervals than we had available from human donations, providing better baseline ultrastructure for detecting any cryopreservation-induced artifacts. However, it is possible that species differences could affect the generalizability of our results. Fourth, we assessed only morphological preservation and did not evaluate biomolecular labeling outcomes such as immunoreactivity, which is the primary rationale for this approach over conventional room temperature or refrigerator storage. Finally, our volumetric assessment used only one type of linear measurement on CT images, which has limited sensitivity for detecting small volume changes and is susceptible to measurement variability due to differences in landmark identification across serial scans. Although this was not the primary focus of our study, future studies would benefit from larger sample sizes and more robust approaches to measuring volume changes, such as serial photography with a visible ruler.

## Conclusions

We present our protocol for long-term storage of aldehyde-fixed whole brains at -20°C using ethylene glycol and sucrose cryoprotection. CT imaging demonstrates that approximately 10 months is required for our protocol to allow for cryoprotectant equilibration in whole human brains, with white matter equilibrating more slowly than grey matter. Our histological validation results, once optimized to allow adequate time for cryoprotectant diffusion into the white matter, corroborates existing data that this protocol maintains brain tissue ultrastructure through the process of cryoprotectant loading, brief storage, and cryoprotectant unloading. Future work should evaluate the long-term biomolecular preservation outcomes and determine whether the protocol can be simplified without compromising tissue quality.

## Abbreviations

ABC: Aldehyde-based cryopreservation
CPA: Cryoprotective agent
CT: Computed tomography
EM: Electron microscopy
H&E: Hematoxylin and Eosin
ICC: Intraclass correlation coefficient
MW: Molecular weight
NBF: Neutral buffered formalin
PBS: Phosphate buffered saline
PMI: Postmortem interval
PVP: Polyvinylpyrrolidone
TEM: Transmission electron microscopy
WSI: Whole slide image

## Author Contributions

**Conceptualization:** M.G., A.K., K.F., J.F.C., A.T.M. **Investigation:** M.G., A.K., A.So., E.L.T., C.D.S. **Formal analysis:** M.G., A.K., A.Sl., A.T.M. **Writing – original draft:** A.T.M. **Writing – review & editing:** M.G., A.K., A.Sl., A.So., E.L.T., C.D.S., K.F., J.F.C., A.T.M.

## Acknowledgements

We would like to acknowledge the Neuropathology Brain Bank & Research CoRE RRID: SCR_027565 at the Icahn School of Medicine at Mount Sinai for their histology and tissue processing services. For the initial protocol, electron microscopy was performed at the Multiscale Microscopy Core, a member of the OHSU University Shared Resource Cores RRID:SCR_009969. For the refined protocol, electron microscopy tissue preparation and imaging were performed at The Microscopy and Advanced Bioimaging CoRE at the Icahn School of Medicine at Mount Sinai. The Icahn School of Medicine at Mount Sinai provided access to library resources.

## Data Availability Statement

Whole slide image and electron microscopy data can be accessed in a public repository on Zenodo, available here: https://zenodo.org/communities/aldehyde_based_cryopreservation. Code and data used for data analysis is available at https://github.com/andymckenzie/Aldehyde_based_cryopreservation.

## Declaration of Generative AI Technologies

In the preparation of this manuscript, the authors used Claude (Anthropic) for both programming assistance and to improve the manuscript’s language. All AI tool-assisted content was reviewed and edited by the authors, who take full responsibility for the final publication.

## Supporting Information

**S1 Data. Serial CT measurements across donor brains**. Maximum brain width measurements for donor IDs #54, #137, and #180 across 11 timepoints during cryoprotectant loading. For each time point, the width measurements were taken at the first coronal slice in which the frontal horns of the lateral ventricles were seen.

## References

1. McKenzie AT, Nnadi O, Slagell KD, Thorn EL, Farrell K, Crary JF. Fluid preservation in brain banking: a review. Free Neuropathol. 2024 Jan;5:5–10. doi:10.17879/freeneuropathology-2024-5373 PubMed PMID: 38690035; PubMed Central PMCID: PMC11058410.

2. McKenzie AT, Thorn EL, Nnadi O, Wróbel B, Kendziorra E, Farrell K, et al. Cryopreservation of brain cell structure: a review. Free Neuropathol. 2024 Jan;5:35. doi:10.17879/freeneuropathology-2024-5883 PubMed PMID: 39844781; PubMed Central PMCID: PMC11753176.

3. de Olmos JS. An improved HRP method for the study of central nervous connections. Exp Brain Res. 1977 Sep 28;29(3–4):541–51. doi:10.1007/BF00236191 PubMed PMID: 72001.

4. Watson REJ, Wiegand SJ, Clough RW, Hoffman GE. Use of cryoprotectant to maintain long-term peptide immunoreactivity and tissue morphology. Peptides. 1986 Feb;7(1):155–9. doi:10.1016/0196-9781(86)90076-8 PubMed PMID: 3520509.

5. Insausti R, Tuñón T, Sobreviela T, Insausti AM, Gonzalo LM. The human entorhinal cortex: a cytoarchitectonic analysis. J Comp Neurol. 1995 May 1;355(2):171–98. doi:10.1002/cne.903550203 PubMed PMID: 7541808.

6. Simmons DM. Cryoprotectant Tissue Storage Solutions: Stability at Lower Temperatures, Longer Storage Times, More Versatile Usage. Microsc Today. 2000 Apr 1;8(3):28. doi:10.1017/S155192950006106X

7. Waldvogel HJ, Curtis MA, Baer K, Rees MI, Faull RLM. Immunohistochemical staining of post-mortem adult human brain sections. Nat Protoc. 2006;1(6):2719–32. doi:10.1038/nprot.2006.354 PubMed PMID: 17406528.

8. Hoffman GE, Le WW. Just cool it! Cryoprotectant anti-freeze in immunocytochemistry and in situ hybridization. Peptides. 2004 Mar;25(3):425–31. doi:10.1016/j.peptides.2004.02.004 PubMed PMID: 15134865.

9. Welikovitch LA, Do Carmo S, Maglóczky Z, Szocsics P, Lőke J, Freund T, et al. Evidence of intraneuronal Aβ accumulation preceding tau pathology in the entorhinal cortex. Acta Neuropathol (Berl). 2018 Dec;136(6):901–17. doi:10.1007/s00401-018-1922-z PubMed PMID: 30362029.

10. Spencer C, Shahidehpour R, Lilek J, Ajroud K, Kawles A, Feldman A, et al. The Use of Sucrose as a Cryoprotectant for Fixed Human Brain Tissue. J Neuropathol Exp Neurol. 2020;79(6):700.

11. Otubo A, Maejima S, Oti T, Satoh K, Ueda Y, Morris JF, et al. Immunoelectron Microscopic Characterization of Vasopressin-Producing Neurons in the Hypothalamo-Pituitary Axis of Non-Human Primates by Use of Formaldehyde-Fixed Tissues Stored at -25 °C for Several Years. Int J Mol Sci. 2021 Aug 25;22(17):9180. doi:10.3390/ijms22179180 PubMed PMID: 34502087; PubMed Central PMCID: PMC8430530.

12. Rosene DL, Roy NJ, Davis BJ. A cryoprotection method that facilitates cutting frozen sections of whole monkey brains for histological and histochemical processing without freezing artifact. J Histochem Cytochem Off J Histochem Soc. 1986 Oct;34(10):1301–15. doi:10.1177/34.10.3745909 PubMed PMID: 3745909.

13. Manger PR, Pillay P, Maseko BC, Bhagwandin A, Gravett N, Moon DJ, et al. Acquisition of brains from the African elephant (Loxodonta africana): perfusion-fixation and dissection. J Neurosci Methods. 2009 Apr 30;179(1):16–21. doi:10.1016/j.jneumeth.2009.01.001 PubMed PMID: 19168095.

14. Manger PR. Collectibles and collections for comparative and evolutionary neurobiological research in Africa. Ann N Y Acad Sci. 2011 May;1225 Suppl 1:E85–93. doi:10.1111/j.1749-6632.2010.05948.x PubMed PMID: 21599700.

15. Kumarasami R, Verma R, Pandurangan K, Ramesh JJ, Pandidurai S, Savoia S, et al. A technology platform for standardized cryoprotection and freezing of large-volume brain tissues for high-resolution histology. Front Neuroanat. 2023;17:1292655. doi:10.3389/fnana.2023.1292655 PubMed PMID: 38020211; PubMed Central PMCID: PMC10651725.

16. Adams JH, Murray MF. Atlas of post-mortem techniques in neuropathology. Cambridge University Press; 1982. doi:10.1017/CBO9780511735479

17. McKee AC. Brain banking: basic science methods. Alzheimer Dis Assoc Disord. 1999;13 Suppl 1:S39–44. PubMed PMID: 10369517.

18. Stempinski ES, Pagano L, Riesterer JL, Adamou SK, Thibault G, Song X, et al. Automated large volume sample preparation for vEM. Methods Cell Biol. 2023;177:1–32. doi:10.1016/bs.mcb.2023.01.009 PubMed PMID: 37451763.

19. Garrood M, Keberle A, Sowa A, Janssen W, Thorn EL, Sanctis CD, et al. Evaluating ultrastructural preservation quality in banked brain tissue. Free Neuropathol. 2025;6:13. doi:10.17879/freeneuropathology-2025-6763 PubMed PMID: 40567433; PubMed Central PMCID: PMC12189018.

20. Koo TK, Li MY. A Guideline of Selecting and Reporting Intraclass Correlation Coefficients for Reliability Research. J Chiropr Med. 2016 Jun;15(2):155–63. doi:10.1016/j.jcm.2016.02.012 PubMed PMID: 27330520; PubMed Central PMCID: PMC4913118.

21. Coleman R, Kogan I. An improved low-formaldehyde embalming fluid to preserve cadavers for anatomy teaching. J Anat. 1998 Apr;192(Pt 3):443–6. doi:10.1046/j.1469-7580.1998.19230443.x PubMed PMID: 9688512; PubMed Central PMCID: PMC1467790.

22. Vlieg RC, Gillespie C, Lee WM. Evaluating passive optical clearing protocols for two-photon deep tissue imaging in adult intact visceral and neuronal organs [Internet]. bioRxiv; 2015 [cited 2024 Nov 22]. p. 018622. Available from: https://www.biorxiv.org/content/10.1101/018622v2 doi:10.1101/018622

23. Krassner MM, Kauffman J, Sowa A, Cialowicz K, Walsh S, Farrell K, et al. Postmortem changes in brain cell structure: a review. Free Neuropathol. 2023 Jan;4:4–10. doi:10.17879/freeneuropathology-2023-4790 PubMed PMID: 37384330; PubMed Central PMCID: PMC10294569.

24. Ethylene Glycol Heat-Transfer Fluid Properties: Density, Data & Charts [Internet]. [cited 2026 Feb 27]. Available from: https://www.engineeringtoolbox.com/ethylene-glycol-d_146.html

25. Drury PJ, Olsen EG, Ross DN. Morphological assessment of sucrose preservation for porcine heart valves. Thorax. 1982 Jun;37(6):466–71. doi:10.1136/thx.37.6.466 PubMed PMID: 7135282; PubMed Central PMCID: PMC459343.

26. Chiffelle TL, Putt FA. Propylene and ethylene glycol as solvents for Sudan IV and Sudan black B. Stain Technol. 1951 Jan;26(1):51–6. doi:10.3109/10520295109113178 PubMed PMID: 14809507.

27. Williamson DI, Russell G. Ethylene glycol as a preservative for marine organisms. Nature. 1965 Jun 26;206(991):1370–1. doi:10.1038/2061370a0 PubMed PMID: 5838256.

28. Young JZ. Osmotic Pressure of Fixing Solutions. Nature. 1935 May;135(3420):823–4. doi:10.1038/135823b0

29. Pallotto M, Watkins PV, Fubara B, Singer JH, Briggman KL. Extracellular space preservation aids the connectomic analysis of neural circuits. eLife. 2015 Dec 9;4:e08206. doi:10.7554/eLife.08206 PubMed PMID: 26650352; PubMed Central PMCID: PMC4764589.

30. Wangensteen D, Bachofen H, Weibel ER. Effects of glutaraldehyde or osmium tetroxide fixation on the osmotic properties of lung cells. J Microsc. 1981 Nov;124(Pt 2):189–96. PubMed PMID: 6798217.

31. Lee RM, McKenzie R, Kobayashi K, Garfield RE, Forrest JB, Daniel EE. Effects of glutaraldehyde fixative osmolarities on smooth muscle cell volume, and osmotic reactivity of the cells after fixation. J Microsc. 1982 Jan;125(Pt 1):77–88. doi:10.1111/j.1365-2818.1982.tb00324.x PubMed PMID: 6806478.

32. Paljärvi L, Garcia JH, Kalimo H. The efficiency of aldehyde fixation for electron microscopy: stabilization of rat brain tissue to withstand osmotic stress. Histochem J. 1979 May;11(3):267–76. doi:10.1007/BF01005026 PubMed PMID: 110731.

33. Carstensen EI, Aldridge WG, Child SZ, Sullivan P, Brown HH. Stability of cells fixed with glutaraldehyde and acrolein. J Cell Biol. 1971 Aug;50(2):529–32. doi:10.1083/jcb.50.2.529 PubMed PMID: 4939525; PubMed Central PMCID: PMC2108264.

34. Penttila A, Kalimo H, Trump BF. Influence of glutaraldehyde and-or osmium tetroxide on cell volume, ion content, mechanical stability, and membrane permeability of Ehrlich ascites tumor cells. J Cell Biol. 1974 Oct;63(1):197–214. doi:10.1083/jcb.63.1.197 PubMed PMID: 4138889; PubMed Central PMCID: PMC2109340.

35. Meissner DH, Schwarz H. Improved cryoprotection and freeze-substitution of embryonic quail retina: a TEM study on ultrastructural preservation. J Electron Microsc Tech. 1990 Apr;14(4):348–56. doi:10.1002/jemt.1060140410 PubMed PMID: 2332811.

36. McIntyre RL, Fahy GM. Aldehyde-stabilized cryopreservation. Cryobiology. 2015 Dec;71(3):448–58. doi:10.1016/j.cryobiol.2015.09.003 PubMed PMID: 26408851.

37. McFadden WC, Walsh H, Richter F, Soudant C, Bryce CH, Hof PR, et al. Perfusion fixation in brain banking: a systematic review. Acta Neuropathol Commun. 2019 Sep 5;7(1):146. doi:10.1186/s40478-019-0799-y PubMed PMID: 31488214; PubMed Central PMCID: PMC6728946.

38. Garrood M, Keberle A, Taylor GA, Thorn EL, Sanctis CD, Farrell K, et al. Mechanical perfusion in brain banking: methods of assessment and relationship to the postmortem interval. Free Neuropathol. 2025;6:20. doi:10.17879/freeneuropathology-2025-8880 PubMed PMID: 41159117; PubMed Central PMCID: PMC12557960.

39. Robards AW, Sleytr UB. Low Temperature Methods in Biological Electron Microscopy. Elsevier Science; 1985. 586 p.

40. Suda I, Kito K, Adachi C. Viability of long term frozen cat brain in vitro. Nature. 1966 Oct 15;212(5059):268–70. doi:10.1038/212268a0 PubMed PMID: 5970120.

41. Suda I, Kito K, Adachi C. Bioelectric discharges of isolated cat brain after revival from years of frozen storage. Brain Res. 1974 Apr 26;70(3):527–31. doi:10.1016/0006-8993(74)90263-7 PubMed PMID: 4821065.

42. van der Valk T, Pečnerová P, Díez-Del-Molino D, Bergström A, Oppenheimer J, Hartmann S, et al. Million-year-old DNA sheds light on the genomic history of mammoths. Nature. 2021 Mar;591(7849):265–9. doi:10.1038/s41586-021-03224-9 PubMed PMID: 33597750; PubMed Central PMCID: PMC7116897.

43. Hattori S, Kiriyama-Tanaka T, Kusubata M, Taga Y, Ebihara T, Kumazawa Y, et al. Preservation of collagen in the soft tissues of frozen mammoths. PloS One. 2021;16(10):e0258699. doi:10.1371/journal.pone.0258699 PubMed PMID: 34714842; PubMed Central PMCID: PMC8555803.

